# Dissecting bacterial resistance and resilience in antibiotic responses

**DOI:** 10.1101/305482

**Authors:** Hannah R. Meredith, Virgile Andreani, Allison J. Lopatkin, Anna J. Lee, Deverick J. Anderson, Gregory Batt, Lingchong You

## Abstract

An essential property of microbial communities is the ability to survive a disturbance. Survival can be achieved through *resistance*, the ability to absorb effects of a disturbance without a significant change, or *resilience*, the ability to recover after being perturbed by a disturbance. These concepts have long been applied to the analysis of ecological systems, though their interpretations are often subject to debate. Here we show that this framework readily lends itself to the dissection of the bacterial response to antibiotic treatment, where both terms can be unambiguously defined. The ability to tolerate the antibiotic treatment in the short term corresponds to resistance, which primarily depends on traits associated with individual cells. In contrast, the ability to recover after being perturbed by an antibiotic corresponds to resilience, which primarily depends on traits associated with the population. This framework effectively reveals the phenotypic signatures of bacterial pathogens expressing extended spectrum β-lactamases (ESBLs), when treated by a β-lactam antibiotic. Our analysis has implications for optimizing treatment of these pathogens using a combination of a β-lactam and a β-lactamase (Bla) inhibitor. In particular, our results underscore the need to dynamically optimize combination treatments based on the quantitative features of the bacterial response to the antibiotic or the Bla inhibitor.

## Introduction

A disturbance is a biological, chemical, or physical event that affects a community^1^. Given that the environment is constantly changing, an essential property of a community is its ability to recover after being disturbed. Responses to a disturbance include resistance, the ability to withstand perturbation in the presence of a disturbance; resilience, the ability to recover after being perturbed by a disturbance; or sensitivity, the inability to withstand or recover from a disturbance^2–5^. Resistance and resilience have been documented in a range of systems and are often determined by different processes^1,2,5,6^. Specifically, resistance is associated with processes that enable the tolerance of, or adaptation to, a disturbance, whereas resilience is associated with recolonization, reproduction, or rapid regrowth^2^. The ability to identify the determinants for resistance versus resilience is crucial for predicting how a given community will respond to a disturbance as well as for designing strategies that will either preserve, change, or eliminate it^3,4,7^. Although resistance and resilience have been defined in the literature for decades^8^, the resistance-resilience framework has not been widely applied. This is partially because it is often difficult to determine the predisturbance state of a population, the definitions vary, and there is a lack of quantitative studies demonstrating how to implement these terms^2,4,9–12^.

Yet, this resistance-resilience framework naturally lends itself to the analysis of bacterial responses to antibiotic treatment. When running susceptibility tests, it is possible to characterize a pre-disturbance state (i.e. no exposure to antibiotic) and there are many ways to quantify the bacterial antibiotic responses (i.e. agar plates, E-test, plate reader, microscopy)^13–15^. Until now, resistance and resilience have not been distinguished in the context of antibiotic responses. Instead, bacteria are classified as resistant if they survive exposure to a set concentration of antibiotic after a set amount of time^16^,^17^ (**Table 1**). However, the apparently similar survival can result from diverse underlying mechanisms^18–22^ that can lead to the ability of individual cells to withstand the treatment or the ability of the population to recover from the initial disturbance. We term the former resistance and the latter resilience.

**Table 1:**
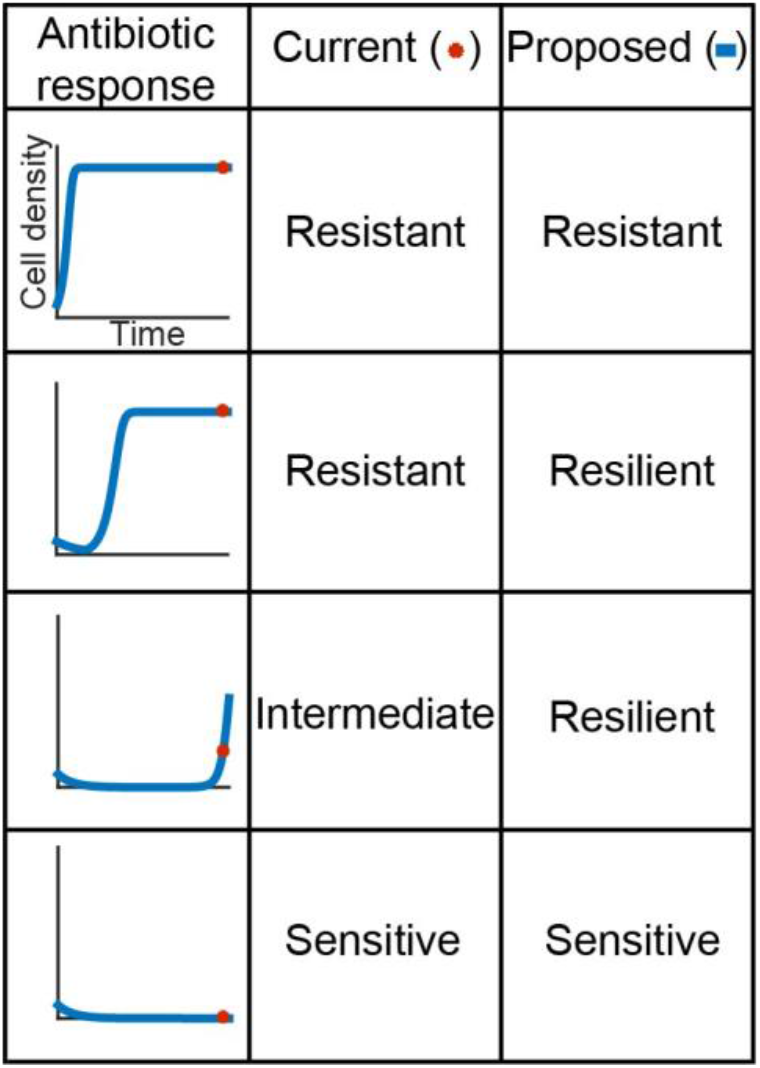
Defining antibiotic responses. Each panel represents a different response to an antibiotic delivered at time = 0. In the first row, the population grows up to carrying capacity, unperturbed. In the second row, the population is perturbed, but eventually recovers in the allotted time. A third population is perturbed and only partially recovers during the allotted time. In the final row, the population is perturbed and does not recover. Currently, antibiotic sensitivity analyses only consider whether bacteria can recover from a set dose of antibiotic in a standard period (red dot). Bacteria that display full recovery are considered resistant, partial recovery are intermediate, and no recovery are sensitive. However, temporal dynamics (blue curve) reveal differences in how a population recovers.

| Antibiotic response | Current (●) | Proposed (▢) |
| --- | --- | --- |
|  | Resistant | Resistant |
|  | Resistant | Resilient |
|  | Intermediate | Resilient |
|  | Sensitive | Sensitive |

Here, we apply these concepts to the analysis of bacterial pathogens producing extended spectrum β-lactamases (ESBLs), which are becoming increasingly prevalent and can degrade many β-lactam antibiotics – the most widely used class of antibiotics in the clinics^23–27^. On one hand, our results offer new insights into the design of antibacterial treatment strategies against one of the fastest-rising types of bacterial pathogens^28–30^. In particular, the resistance-resilience framework reveals the phenotypic signatures of different ESBL-producing bacteria and underscores the critical need to implement adjustable formulations of combination treatments. On the other, the framework we illustrate here is generally applicable for the analysis of bacterial populations to any environmental perturbations.

## Results

### Temporal dynamics of ESBL-producing bacteria in response to β-lactam treatment

The dynamics of an ESBL-producing population are uniquely suited for illustrating resistance and resilience during disturbances. In the absence of antibiotic treatment, the population grows approximately exponentially before reaching the stationary phase, upon depletion of nutrients (**Figure 1A**). Thanks to the expression of a β-lactamase (Bla) anchored in their periplasm, these bacteria can degrade antibiotic that diffuses across the outer membrane^31^. However, if Bla expression is moderate, these bacteria can still be lysed by a β-lactam antibiotic at a sufficiently high concentration (**Figure 1B**). As this occurs, Bla is released into the environment due to membrane leakage (from a cell not-yet lysed) or cell lysis^32^. If sufficient Bla (periplasmic and extracellular) is present, the antibiotic is degraded in time for the population to recover before all cells are lysed. A population’s recovery depends on collective tolerance (**Supplementary Fig. 1**): it must have a sufficiently high density when treated so enough Bla is collectively produced to remove the antibiotic before all bacteria are lysed. If the initial density is too low, then insufficient Bla will be produced to protect the population from antibiotic exposure.

**Figure 1:**
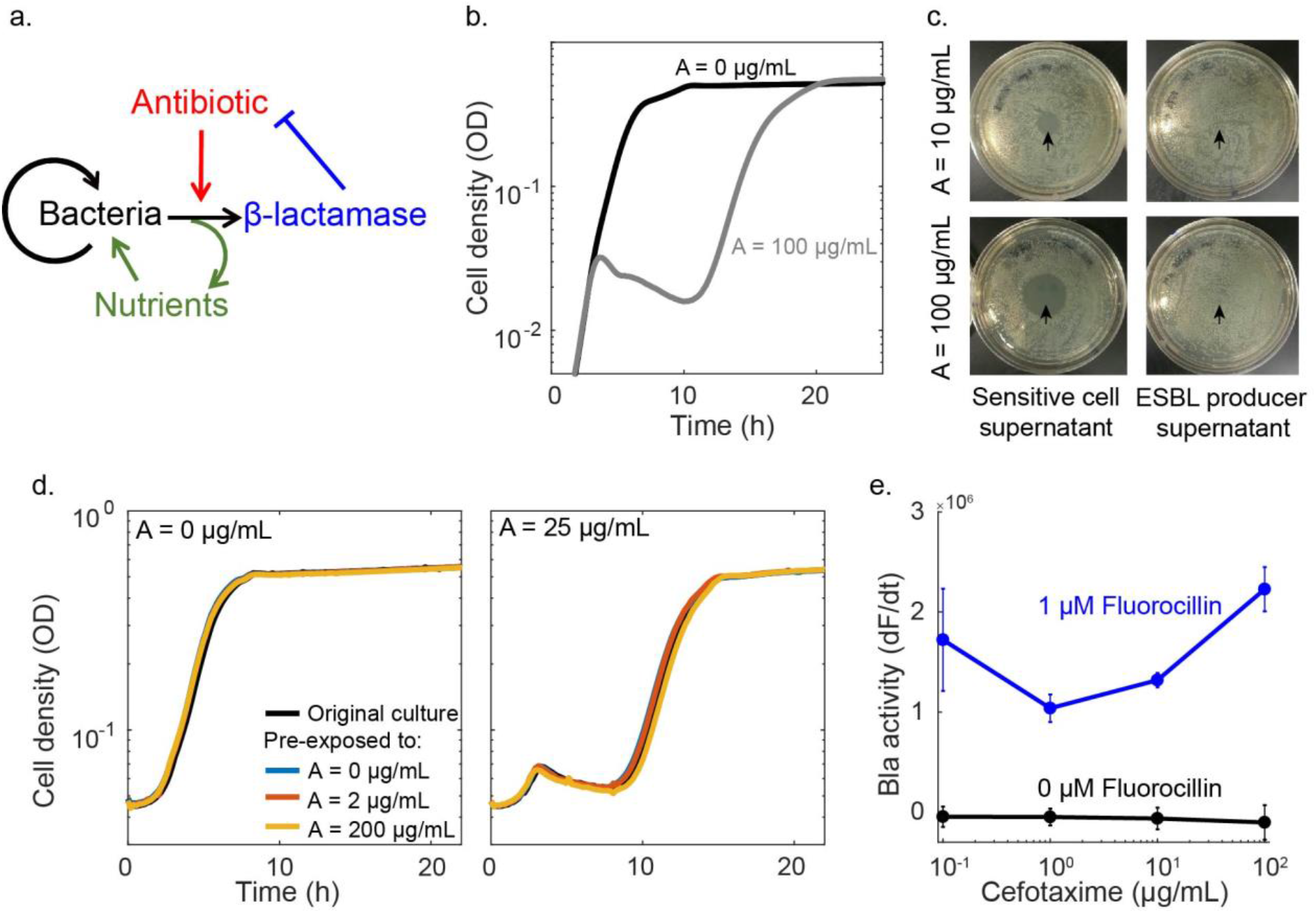
Response of an ESBL-producing population to cefotaxime, a β-lactam. **a. Schematic of antibiotic response of an ESBL-producing population.** In the absence of antibiotic, bacteria reproduce and consume nutrients. Upon the introduction of an antibiotic, some of the bacteria undergo lysis and release β-lactamase and a small amount of recyclable nutrients into the environment. The released Bla degrades the antibiotic (blue inhibition arm). **b. Time course of antibiotic response.** In the absence of an antibiotic (black curve), the bacteria grow to carrying capacity without any delay. In the presence of sufficient antibiotic (grey curve; A=100 μg/mL cefotaxime), the population displays the characteristic crash, as the cells lyse, and recovery after the Bla released from lysed cells degrades the antibiotic. **c. ESBL significantly degraded cefotaxime in a short time window.** The supernatant from a culture of sensitive cells still contained significant concentrations of cefotaxime, as depicted by the zones of inhibition in the lawn of sensitive cells (strain MC4100, left column). The supernatant from the culture containing ESBL producing bacteria did not contain significant concentrations of cefotaxime, as depicted by the full lawns (right column). Arrowheads indicate where supernatant was placed on the agar plate. **d. Populations previously exposed to cefotaxime exhibited the same temporal dynamics.** Culture was treated with a range of antibiotic concentrations. After 24 hours, bacteria from the recovered population were used to re-inoculate fresh media with or without 25μg/mL cefotaxime. During the 2^nd^ round of treatment, the time courses from the populations previously exposed to 0, 2, or 200 μg/mL cefotaxime were identical, suggesting that the population recovery was unlikely due to mutants or phenotypic variants with increased tolerance. **e. Bla production is not induced by cefotaxime.** We used Fluorocillin to determine that the isolate’s Bla production is not significantly increased by the addition of antibiotic. Here, the Bla activity present in a population after 3 hours of exposure to a range of antibiotic concentrations was quantified by the rate at which Fluorocillin was hydrolyzed and produced green fluorescence. One way ANOVAs indicate that the increase in fluorescence recorded was insignificant when compared to the control.

Here, we showed that the population recovery was indeed due to the Bla degrading the antibiotic, indicated by the level of antibiotic activity in the supernatant after 6 hours of exposure (**Figure 1C**). Sensitive bacteria do not produce Bla and cannot break down the antibiotic, thus the antibiotic remaining in the supernatant could inhibit the growth of sensitive bacteria. At a higher initial antibiotic concentration, the same amount of supernatant generated a larger zone of inhibition. In contrast, the bacteria producing ESBLs sufficiently degraded the antibiotic at both doses during the incubation period, as evidenced by the inability of the resulting supernatants to inhibit growth of sensitive cells.

To test if the population recovery was due to the selection of a more resistant or resilient subpopulation, we collected ESBL producing bacteria that had recovered from an antibiotic treatment and re-exposed them to a range of antibiotic concentrations. The resulting antibiotic responses were similar, regardless of previous exposure concentrations (**Figure 1D**). This shows that the recovery was not due to the selection of a sub-population with enhanced tolerance, which is consistent with the notion of antibiotic degradation due to the Bla released from cell lysis.

We also tested if the antibiotic induced the production of Bla by using Fluorocillin, a substrate that fluoresces green when degraded by Bla, allowing for the real-time visualization of Bla activity. After incubating the isolate with different concentrations of cefotaxime for 3 hours, we added Fluorocillin to the sonicated culture to quantify the total Bla present. There was no significant increase in fluorescence as a function of antibiotic concentration (**Figure 1E**) at the p < 0.05 level [F(1, 6) = 2.44, p = 0.17] and [F(1, 6) = 3.31, p = 0.12] for A = 10 and 100 μg/mL, respectively. There was a slight, but significant, decrease in fluorescence for A= 1 μg/mL [F(1, 6) = 6.68, p = 0.04]. Overall, Bla production is not induced by exposure to this range of antibiotic concentrations.

### Defining resistance and resilience

A population can survive a disturbance due to its resistance or resilience^1,2,9^. In general, resistance refers to the ability of a population (or a community) to *withstand* the disturbance, whereas resilience refers to the ability to *recover* after suffering from the disturbance. Both properties are evident in the temporal dynamics of an ESBL-producing population in response to β-lactam treatment (**Figure 1**). Here, we quantify resistance as the ability of the population to not deviate from the pre-antibiotic-treatment state (as quantified by the growth rate). Mathematically, we define resistance at the time of maximum change in a treated population’s net growth rate (ρ_A_) normalized by the untreated population’s net growth rate at the same point in time (ρ_0_). When dealing with the experimental data, our analysis accounts for the time delay in lysis caused by a β-lactam^33^.

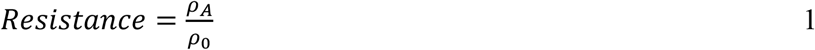

By this definition, resistance primarily reflects the instantaneous response of individual cells, but manifested at the population level. In particular, the magnitude of the metric depends on the probability by which an average bacterium is lysed by the antibiotic. This probability is directly determined by the expression level and activity of Bla in the bacterium, as well as the extracellular concentration of the antibiotic. In our analysis, we use optical density (OD) as a measure of the cell density.

We quantify resilience as the rate of recovery by the population after experiencing the initial crash (**Figure 1**), by using the time needed for a population to reach 50% of its carrying capacity (*T*^50%^). With increasing antibiotic concentration, more cells will lyse in the process of degrading the antibiotic, thus increasing the resulting recovery time. The more resilient a population is, the faster it can return to a normal state after being perturbed by an antibiotic. We define resilience as the inverse of the treated population’s 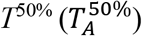, normalized by the untreated population’s 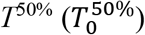 (Figure 2A-B, **Supplementary Fig. 2**)^1^. The inverse is taken to reflect the fact that increasing the recovery time corresponds to a decrease in resilience.

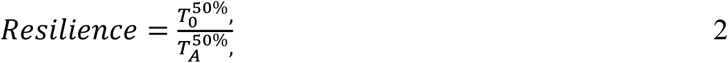

**Figure 2:**
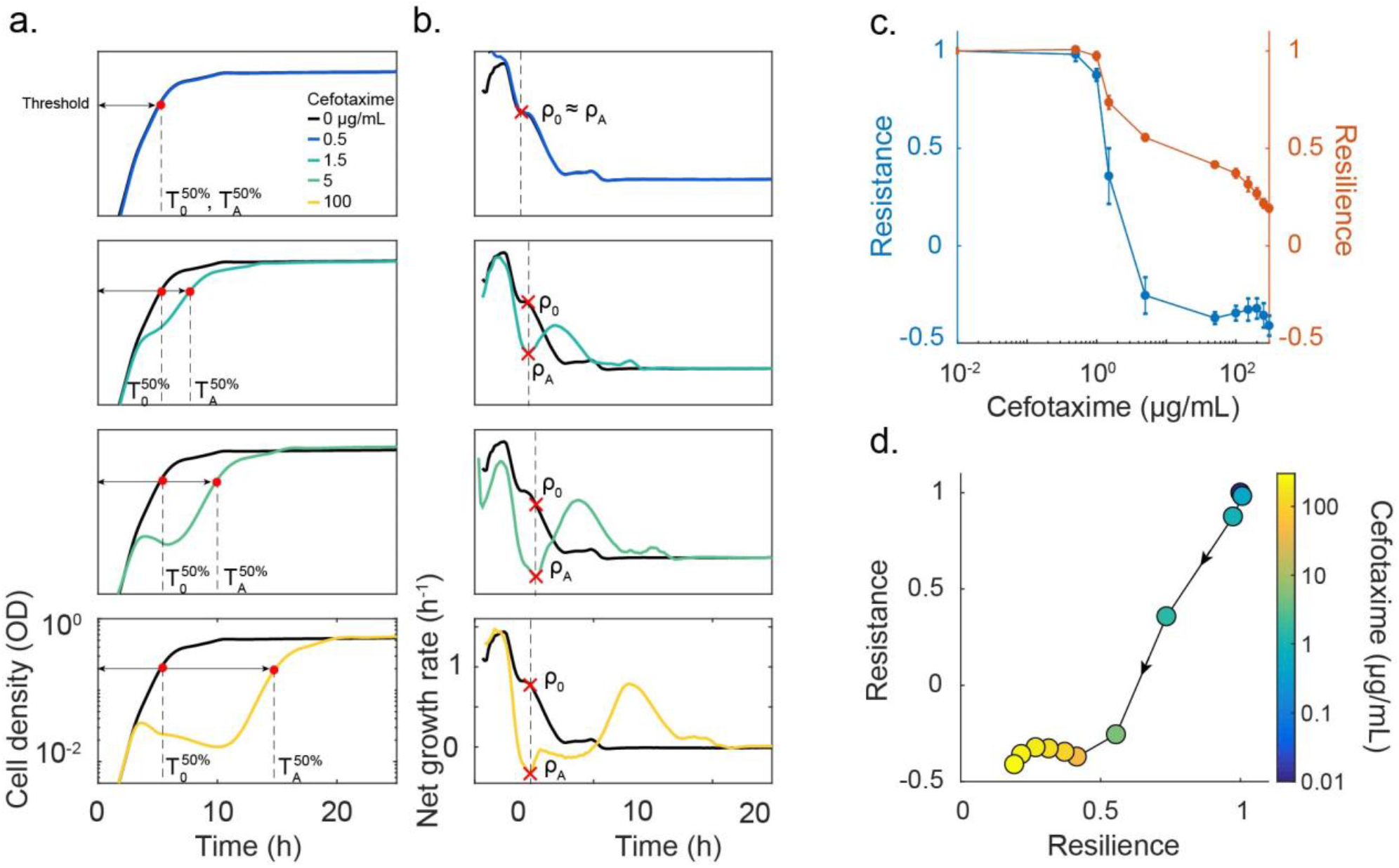
Quantifying resistance and resilience. **(A) Time courses are used to quantify a population’s resilience.** When no antibiotic was added (black curve), the population grew up unperturbed and reached a target threshold density (here 50% of the carrying capacity) in time = *T*^50%^. If the antibiotic concentration added was very low (blue curve, A=0.5 μg/mL cefotaxime), then the population reached the threshold density in a similar time to the untreated population. As the antibiotic concentration increased, the degree of lysis increased and affected the time necessary for the treated population to reach the threshold 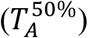. We characterized the population’s resilience for a range of antibiotic concentrations as the inverse ratio of the times to the half maximal carrying capacity 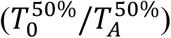. **(B) Net growth rate quantifies population’s resistance.** When no antibiotic was added, the population’s net growth rate decreased over time as it consumed the available nutrients and approached stationary phase. When a low dose of antibiotic was added, the net growth was not significantly altered (blue curve). In this instance, the treated population’s maximum net growth rate is recorded as ρ_A_ and compared to the untreated population’s net growth rate at the same time (ρ_0_). As the antibiotic concentration increased, the net growth rate curve of the treated population deviated more from the untreated curve. For each antibiotic concentration, ρ_A_ was recorded as the net growth rate at the point of maximum negative deviation from the untreated population and normalized by ρ_0_. We characterized a population’s resistance as the ratio of recovery times (ρ_A_/ρ_0_). **(C) Resistance and resilience as functions of the cefotaxime concentration.** At low doses of cefotaxime, the population was resistant and resilient, showing little disturbance after exposure (see Fig. 2A-B). As the antibiotic concentration was increased, resistance and resilience decreased due to the increase in cell lysis causing the net growth to decrease and the time to the half max density increased. Once the population underwent a crash, the resistance was minimized and resilience became the dominating factor for survival. **(D) The resistance-resilience map defines a phenotypic signature.** Using the same data as in Fig. 2A-C, the resistance-resilience framework can visualize the shift in a population’s antibiotic response. When the antibiotic concentration was 0 or very low, the population’s response displayed high resistance and resilience. Once the antibiotic concentration increased to 5 μg/mL, the population’s response shifted to a position where resistance was minimized and resilience dominated the antibiotic response. With further increase in antibiotic concentration, the resistance level continued to decrease. An effective treatment should minimize both resistance and resilience. Dot color references the antibiotic concentration used at that point, arrows indicate direction of increasing antibiotic concentration.

By this definition, resilience reflects a long-term response and primarily depends on population-level traits: when a single bacterium can no longer survive the effects of antibiotic, the population is initially impacted; however, collective antibiotic tolerance can allow for the population to outlast the disturbance and recover after being perturbed.

For each bacterial strain, the degree of resistance or resilience depends on the type and dose of antibiotic used. At low antibiotic concentrations, the population experiences little or no disturbance and thus is characterized with relatively high resistance and resilience (**Figure 2C**). At intermediate concentrations (A=1.5 μg/mL), the population’s recovery displays a decline in both resistance and resilience because the antibiotic concentration is high enough to induce some lysis and slow the net growth rate and delay the recovery time. Once the antibiotic concentration is high enough to induce a population crash (A>5 μg/mL), the recovery of the population shifts to being dominated by resilience. If the antibiotic concentration exceeds the threshold for population recovery, the resulting resilience and resistance will be minimal. This resistance-resilience framework effectively reveals the phenotypic signature of each strain (**Figure 2D**) when treated by a β-lactam.

In our experiments, the OD values are sufficiently small such that they are proportional to the total biomass^33^. Debris from lysed cells can also contribute to the OD values. However, this contribution is negligible except for OD values that are near the baseline (when cells are lysed). As such, it has negligible impact on calculated values of resistance or resilience.

### Determinants of resistance and resilience

To help further our investigation, we developed a kinetic model to describe the temporal response of a bacterial population constitutively expressing Bla to a β-lactam antibiotic. When no antibiotic is present, the population grows to carrying capacity without delay; however, once the antibiotic concentration is high enough, the population density undergoes a crash as a significant portion of the population is lysed by the antibiotic, and a recovery, as the released Bla degrades the antibiotic (**Figure 3A, Supplementary Fig. 3**). We chose to simplify the system by lumping the activity of intra- and extra-cellular Bla, based on direct measurements that suggest that extracellular Bla plays a much greater role once significant lysis has occurred (**Supplementary Fig. 4**).

**Figure 3.**
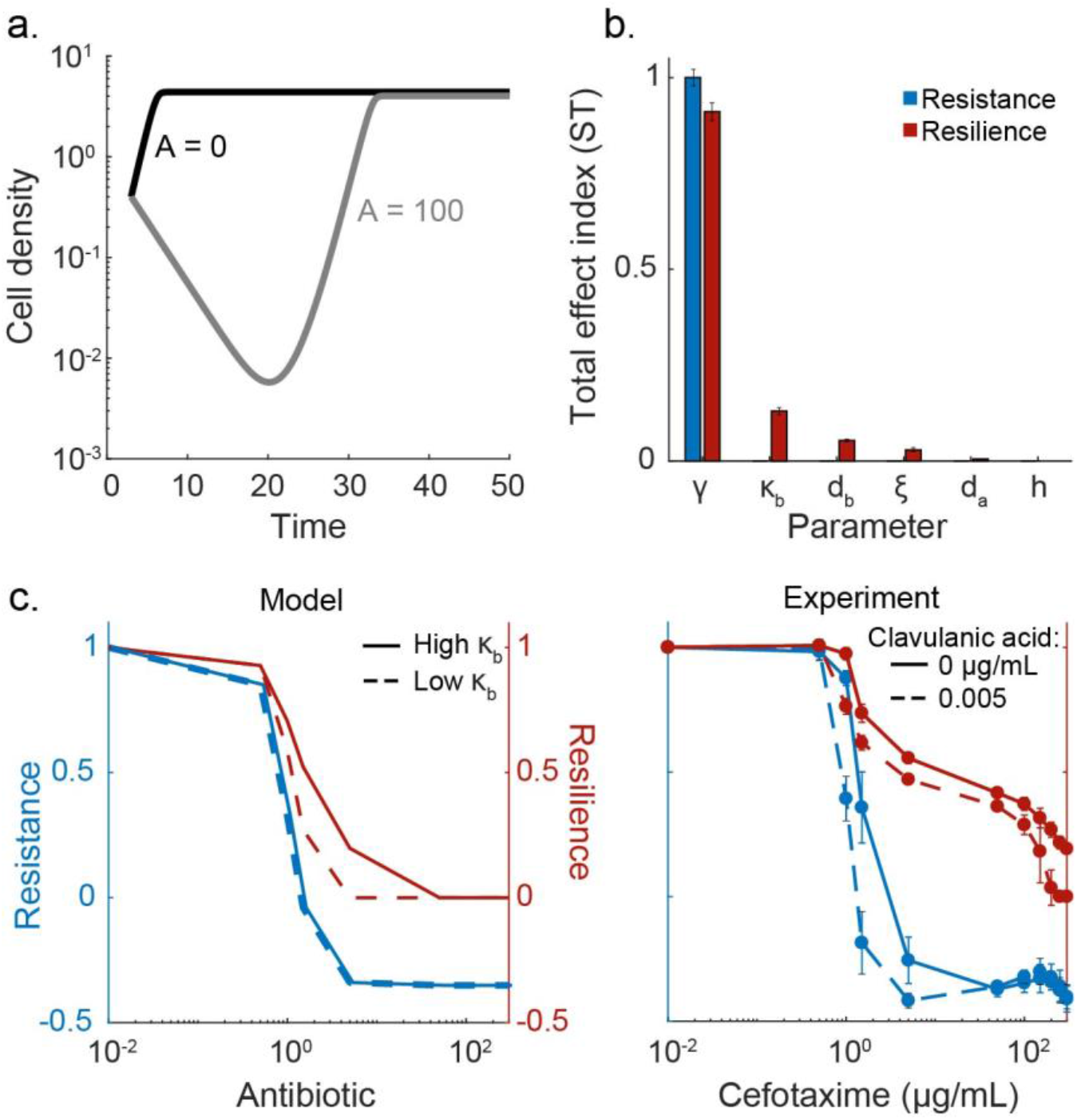
Modeling reveals key determinants of resistance and resilience. **(A) Simulated time courses of an ESBL-producing isolate with and without an antibiotic.** The characteristic “crash and recovery” is generated once the antibiotic concentration is high enough. **(B) Sensitivity analysis reveals determinants of resistance and resilience.** Total effect indices (ST) for resistance and resilience are reported for each parameter (A = 100 μg/mL). Resistance is most affected by the lysis rate (γ). The remaining parameters did not significantly affect the system’s resistance, but did affect resilience. The most influential parameters included the maximum lysis rate (*γ*), Bla activity (*κ_b_*), the turnover rate of Bla (*d_b_*), and the amount of nutrients released during lysis (*ξ*). **(C) Modulating resistance and resilience by tuning Bla activity.** We altered Bla activity in the model (left column) or experimentally added clavulanic acid (right column) in combination with a range of antibiotic concentrations. Here, a low Bla activity corresponds to *κ_b_* = 0 in the model or 0.005 μg/mL clavulanic acid in the experiment. A high Bla activity corresponds to *κ_b_* = 0.35 in the model or no clavulanic acid in the experiment. Reducing Bla activity increased the population’s sensitivity, causing both resistance and resilience to decrease at lower concentrations of antibiotic.

Global sensitivity analysis was used to determine which parameters influenced resistance and resilience under a range of antibiotic concentrations. Briefly, the Sobol method calculates resilience and resistance for a range of parameter values and breaks down the variation for each into fractions that can be attributed to one or more parameters^34^. Here, we reported the total effect index, ST, which reflects how much a parameter and all its interactions with any other parameters contributes to the variation in resistance and resilience at a particular antibiotic concentration (**Figure 3B, Supplementary Fig. 5**). Comparing the ST for each parameter when A=100 μg/mL reveals that resistance is only sensitive to the maximum lysis rate (*ST_γ_* = 1±0.02). The sensitivity analysis revealed that all parameters affect resilience to varying degrees, depending on the antibiotic concentration. Resilience is sensitive to the maximum lysis rate (*ST_γ_*= 0.9 ± 0.02), Bla activity 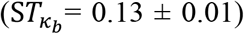, the turnover rate of Bla 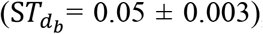, and the amount of nutrients recycled from cell lysis (*ST_ξ_* = 0.03 ± 0.01). These parameters determine the collective ability of the population to remove the antibiotic, underscoring the notion that resilience is a population-level trait.

We tested the influence of Bla activity computationally by varying *κ_b_*, and experimentally by using clavulanic acid, a well-characterized Bla inhibitor (**Figure 3C, Supplementary Fig. 6**). With increased Bla inhibition, the population became more sensitive to the antibiotic, thus resulting in the antibiotic response shifting from relying on both resistance and resilience to just resilience at a lower concentration of antibiotic. Furthermore, the population with significantly reduced Bla activity did not survive exposure to the higher concentrations of antibiotic (resistance and resilience were both reduced).

### Phenotypic signatures of bacterial responses in the resistance-resilience framework

Given that Bla inhibitors are commonly used to restore sensitivity to some β-lactam antibiotics^31^, we explored the implications of using the resistance-resilience framework to optimize combination treatments by analyzing the response of four different ESBL-producing isolates (**Supplementary Table 1**). We exposed each isolate to different combinations of antibiotic and Bla inhibitor concentrations and recorded their antibiotic responses. The resistance and resilience for each scenario were calculated and plotted against each other (**Figure 4A**). The framework revealed how a small dose of antibiotic (5 μg/mL cefotaxime) could minimize resistance for Isolate I, but larger doses were needed to minimize resilience. However, resilience could be minimized with a small dose of Bla inhibitor (0.05 μg/mL clavulanic acid), when in combination with the antibiotic. A similar trend was observed in Isolates II and IV, with treatments using higher concentrations of Bla inhibitor requiring less antibiotic to minimize resistance and resilience. Isolate III, however, was not affected by increasing concentrations of Bla inhibitor, as seen by the overlapping resistance-resilience curves. Only the highest concentration of Bla inhibitor (0.5 μg/mL) in combination with a high concentration of antibiotic (150 μg/mL) prevented the population from recovering.

**Figure 4.**
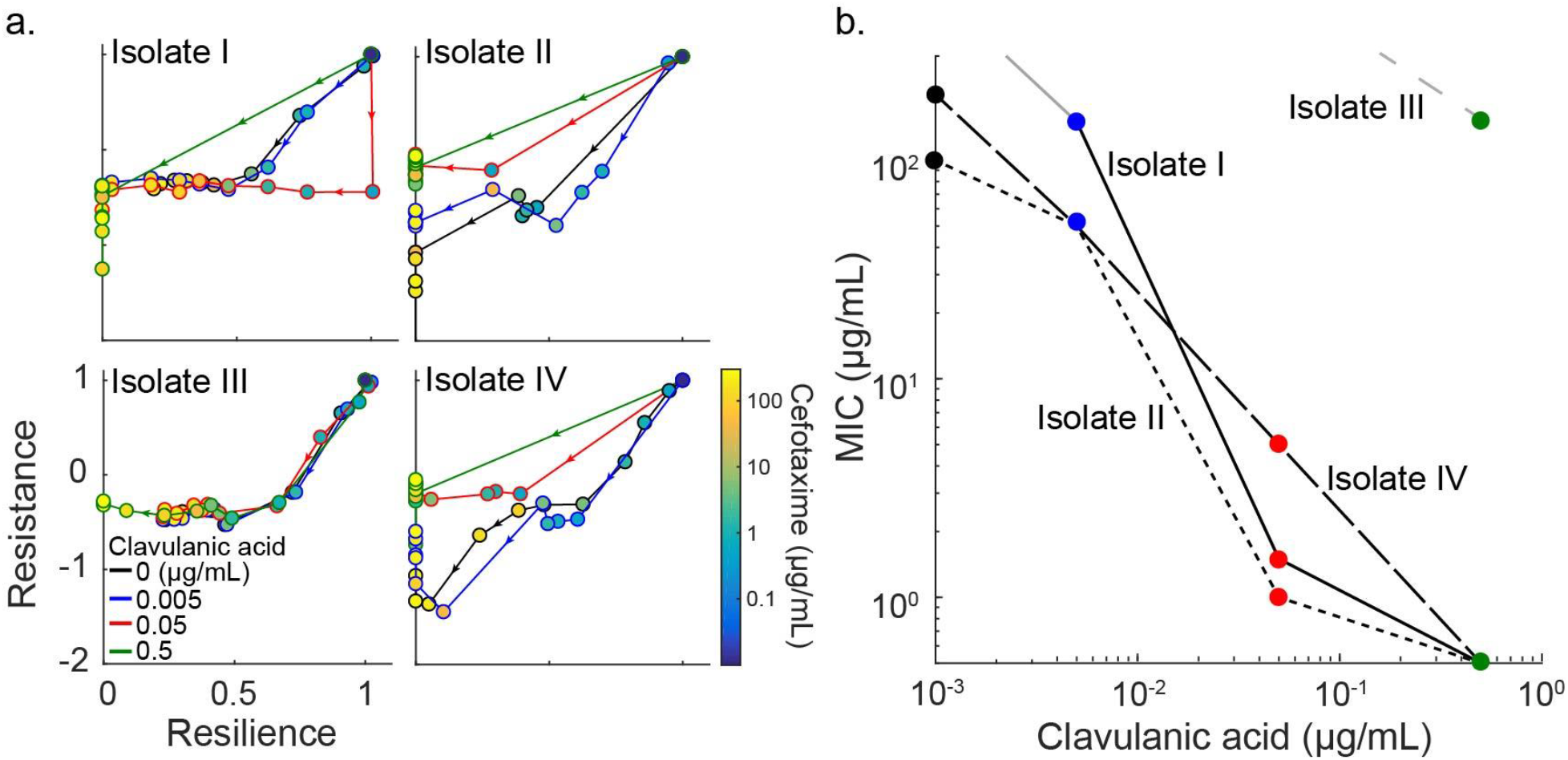
Diverse phenotypic responses by different ESBL-producing isolates. **(A) Responses to combinations of Bla inhibitor and antibiotic concentrations.** In general, with increasing antibiotic concentrations, the population responses start out as both resistant and resilient (top right of each subplot). The resistance is minimized with low doses of antibiotic, giving way to a response driven by resilience. As antibiotic concentration increased, the corresponding resilience decreased. With the addition of clavulanic acid, the concentration at which the response lost its resistance and then resilience is lowered. For Isolates I, II and IV, the highest clavulanic acid concentration (0.5 μg/mL, green curve) causes the antibiotic response to shift directly from resistant and resilient to sensitive with the lowest dose of cefotaxime (0.5 μg/mL). Isolate III was not as affected by clavulanic acid. Color of the curves indicates concentration of clavulanic acid. Color of the dots indicates concentration of cefotaxime and arrowheads indicate direction of increasing antibiotic concentration. **(B) Dependence of MIC on Bla inhibition.** For each ESBL-producing isolate, the MIC corresponds to the lowest concentration of cefotaxime needed to prevent recovery in 48 hours. One isolate, Isolate III, was not very susceptible to clavulanic acid, suggesting this intervention method would not be optimal in this case. Here, different line patterns represent the isolate and the symbol color represents the concentration of clavulanic acid, as defined in (Fig. 4A). The greyed lines indicate that the population recovered under all tested concentrations of cefotaxime tested at that concentration of clavulanic acid, therefore the MIC could not be calculated (Isolate I: clavulanic acid < 0.005 μg/mL; Isolate IV: clavulanic acid < 0.5 μg/mL).

Using the resistance-resilience framework, we determined that the minimum inhibitory concentration (MIC) of antibiotic for each level of Bla inhibitor was unique for each isolate (**Figure 4B**). Here, the MIC is defined as the concentration necessary to prevent the population from recovering within 48 hours of being exposed to treatment. For example, when 0.05 μg/mL clavulanic acid was used, the MIC was 5, 1.5, 1 and > 300 μg/mL cefotaxime for Isolates I, II, III, and IV, respectively. This diversity in the treatment responses can be explained by the expression of different or additional types of Bla that exhibit different sensitivity to inhibition by clavulanic acid. For instance, isolates producing cephalosporinases or chromosomally mediated Bla have been shown to be poorly inhibited by clavulanic acid^31,35,36^. These results underscore a critical caveat in using predetermined formulations of β-lactam/Bla inhibitor combinations to combat ESBL-producing pathogens, which is currently a standard practice^31,37,38^. Given the diversity of the phenotypic responses by the different isolates, quantitative measurements on how a strain responds to an antibiotic and a Bla inhibitor are necessary to predict the outcome of a particular combination treatment.

## Discussion

Resistance and resilience provide a powerful framework to dissect the contributions of different factors underlying a population’s response to a disturbance. Despite their appeal, resistance and resilience have had limited applications due to differing definitions^1,2^ and a lack of quantitative studies. Our analysis of an ESBL-producing clinical isolate’s response to a β-lactam antibiotic serves as a concrete example that illustrates the dichotomy between resistance associated with single cells and resilience associated with populations. That is, resilience can be considered a cooperative trait by a group of bacteria (clonal or mixed).

Our work provides a concrete procedure to quantify resistance and resilience in a population (clonal or mixed) in response to neutral or negative perturbations, as both metrics can be uniquely defined from the time course of the population. As such, it can be applied to diverse situations. For example, some bacteria are resistant to xeric stress due to the disturbance triggering their adaptive mechanisms^39^. Specifically, a xerotolerant cell survives a dry spell by decreasing its energy consumption, protecting its DNA from damaging reactive oxygen species, stabilizing its membrane, and preventing intracellular water loss. Another example of resistance being a single cell level response is the production of heat-shock proteins to enable the cell’s survival of stressful conditions, such as extreme temperatures^40,41^. As for resilience, a population of cyanobacteria has been shown to depend on its density to survive high levels of light that are damaging to single cells. Mutual shading is a density dependent phenomenon achieved when the damaged cyanobacteria that are closer to the light source provide shade to their lower neighbors, thus allowing the population to regrow in lower, less damaging levels of light^42,43^. Additionally, the microbiome is resilient to diet changes, antibiotic exposure and invasion by new species due to population level attributes such as species richness and function response diversity^44^.

In our analysis, the framework is applied to the dynamics of a homogenous population, where the resistance and resilience both depend on the average properties of cells. However, this framework is also applicable for dissecting bacterial responses to antibiotics for heterogeneous populations (**Supplementary Fig. 7**). In the extreme case, a population’s antibiotic response can be driven by a small subpopulation of persisters, or slow-growing or dormant bacteria that are not sensitive to antibiotics^15,45^. Upon antibiotic exposure, most of the population is killed, leaving the persisters behind. Once the antibiotic has been removed, the persisters, which are genetically identical to their sensitive counterparts, spontaneously switch back into the normal, growing state and reestablish the colony. This antibiotic response would be characterized with low resistance and high resilience due to the presence of persisters.

Alternatively, bacteria that have mutated forms of a given antibiotic’s target are resistant to that antibiotic^46^. For instance, vancomycin-resistant *Enterococcus faecalis* (VRE) has mutated the end of a peptidoglycan strand, a component necessary for cell wall synthesis and the target for vancomycin^47^. This mutation reduces the peptidoglycan’s binding affinity for vancomycin by 1000fold. Because a single bacterium of VRE has this mutation and ability to withstand much higher antibiotic concentrations, an entire population’s antibiotic response would be characterized as high resistance.

Our framework is connected to another in quantifying bacterial responses. In particular, Artemova and colleagues introduced the concept of single-cell minimal inhibitory concentration (scMIC)^48^ to describe the susceptibility of individual bacteria to a β-lactam antibiotic^33^. This concept is in contrast to the concept of minimal inhibitory concentration (MIC), which is determined as the collective response of a population. The scMIC concept operates at the same level as resistance described in our study. Both reflect the ability of individual cells to survive an antibiotic except that resistance reflects the ability of an average cell. As a result, both depend on parameters associated with individual cells, including the maximum lysis rate. In contrast, resilience and MIC result from the collective behavior of the entire population. Both also depend on the inoculum size of the population, but resilience has the advantage of accounting for the temporal dynamics.

In addition to dissecting an antibiotic response into its components and corresponding attributes, the resistance-resilience framework has demonstrated the need to modulate combination treatments. Comparing the resistance-resilience fingerprint of different isolates under varied concentrations of cefotaxime and clavulanic acid revealed how varied their responses are to Bla inhibition. As one way to extend the efficacy of a β-lactam is by pairing it with a Bla inhibitor^49–51^, this observation is relevant for guiding the optimization of combination treatments. Currently, there are a few versions of the treatment combining clavulanic acid and amoxicillin for clinical use; however, the concentration of clavulanic acid is kept constant between the versions while the amoxicillin concentration is changed^52–54^. Early studies suggest this clavulanic acid concentration was selected to minimize patient side effects and maintain a sufficiently high serum concentration of clavulanic acid^31,37,38^. Nevertheless, our finding suggests that the clavulanic acid concentration can be optimized within a safe range to reduce the amount of antibiotic necessary and minimize the resistance and resilience of a given isolate. Furthermore, a recent study found that different ratios of inhibitor to antibiotic could influence the rate and mechanism of antibiotic resistance that bacteria develop^55^.

## Methods

1. *Bacterial strains, growth media, and culturing conditions:* We characterized bacterial isolates from a library assembled by the Duke Hospital Infectious Disease department. This library consists of approximately 80 isolates that have been identified as ESBL producers. Unless otherwise noted, *K. pneumoniae* isolate DICON 005 was used. As a sensitive bacteria control, *E. coli* MC4100 cells were used. Unless otherwise indicated, experiments were conducted in M9 medium [1× M9 salts (48 mM Na2HPO4, 22 mM KH2PO4, 862 mM NaCl, 19 mM NH4Cl), 0.4% glucose, 0.2% casamino acids (Teknova), 0.5% thiamine (Sigma), 2 mM MgSO4, 0.1 mM CaCl2]. For overnight cultures, we inoculated single colonies from an agar plate into 2 mL M9 and incubated them for 12 hours at 30°C.
2. *Measuring time courses:* (**Supplementary Fig. 8**) 1 mL overnight culture was washed (centrifuged for 5 minutes at 13000rpm, discarded supernatant, and resuspended in 1 mL fresh M9) and the optical density (OD) was adjusted to 0.5 OD_600_ by adding the appropriate volume of M9. The culture was then diluted 1000fold in fresh M9 and the appropriate amount of cefotaxime (Sigma) was added to achieve a range of concentrations from 0-300 μg/mL. A 96-well plate (Costar) was loaded with 200 μL of culture per well and topped with 50 μL mineral oil (Sigma) to prevent evaporation. The plate was loaded into a Tecan Infinite M200 PRO microplate reader (chamber temperature maintained at 30°C) and OD_600_ readings were measured every 10 minutes for 48 hours with intermittent plate shaking. Unless otherwise noted, each condition tested consisted of 4 technical replicates that, when averaged, did not need to include error bars.
3. *Cefotaxime activity level:* (**Supplementary Fig. 9**) ESBL-producing Isolate I and sensitive strain MC4100 were cultured in M9 for 12 hours at 30°C. One mL of each overnight culture was washed (centrifuged for 5 minutes at 13000rpm, discarded supernatant, and resuspended in 1 mL fresh M9) and the OD was adjusted to 0.5 OD_600_ by adding the appropriate volume of M9. The culture was then diluted 1000fold in 4 mL of fresh M9 and the appropriate amount of cefotaxime was added to achieve final concentrations of 0, 10 or 100 μg/mL. The cultures were incubated at 37°C for 6 hours. At this time, lawns of sensitive cells were prepared by spreading 5 μL of the 1000fold diluted MC4100 culture onto agar plates. Supernatant from the cultures incubated with cefotaxime were prepared by spinning down 0.5 mL culture with 5 μg/mL clavulanic acid (to prevent further Bla activity). 4 μL of the supernatant were dropped into the center of the agar plates, which were then incubated for 16 hours at 37°C. The zones of inhibition were recorded by camera.
4. *Selective pressure:* (**Supplementary Fig. 10**) After conducting a 24-hour time course, as described in (2), culture that had been exposed to 0, 2 ug/mL, and 200 ug/mL of cefotaxime were diluted 10fold in fresh M9 (antibiotic free) and incubated at 37°C for 3 hours. The recovered cultures were then used to run another time course using the same antibiotic concentrations used in the previous round of treatment.
5. *Quantifying Bla activity:* 1 mL overnight culture was washed (centrifuged for 5 minutes at 13000rpm, discarded supernatant, and resuspended in 1 mL fresh M9) and diluted 1000x in fresh M9. The appropriate amount of cefotaxime (Sigma) was added to achieve 0, 1, 10, and 100 μg/mL. The cultures were incubated for 3 hours at 30°C. For each culture, 1 mL was kept on ice to preserve the population density at the 3 hour time point. 2 mL were spun down (5 minutes, 13000rpm) and resuspended in 2 mL water before being sonicated (20 amp, duration 1 minute in 4°C room) to release periplasmic Bla. The sonicated culture was diluted 10fold in water and then treated with 0 or 1 uM of Fluorocillin. A 96-well plate was loaded with 200 μL of each culture (whole cells), sonicate with and without Fluorocillin, and topped with 50 μL mineral oil. The plate was loaded into a Tecan Infinite M200 PRO microplate reader (chamber temperature maintained at 25°C) and OD_600_ and GFP readings were measured every 10 minutes for 1.5 hours. The GFP measurements of the sonicated samples were plotted over time and the slope was calculated. The slope was normalized by the relevant culture’s OD measurement. A one-way ANOVA was used to determine any significant differences between the conditions.
6. *Quantifying internal vs external Bla activity*: ESBL producing isolates were incubated with 0, 0.01, 1, 10 and 100 μ/mL cefotaxime for 12 hours at 30°C. At that point, the culture was centrifuged (13000rpm, 5 minutes) to separate the supernatant and whole cells. The supernatant was removed and the whole cells resuspended in fresh M9. A portion of the whole cells was removed and sonicated (20 amp, duration 1 minute in 4°C room) to release Bla from the periplasmic space. Bla activity in each component (supernatant, sonicated whole cells, and whole cells) was quantified by adding 1 μM Fluorocillin, and monitoring the change in green fluorescence with a plate reader. A 96-well plate was loaded with 200 μL of each mixture and topped with 50 μL mineral oil. The plate was loaded into a Tecan Infinite M200 PRO microplate reader (chamber temperature maintained at 25°C) and OD600 and GFP readings were measured every 10 minutes for 1.5 hours. The GFP measurements of the samples were plotted over time and the slope was calculated. The slope was normalized by the whole cell’s OD measurement.
7. *Varying Bla inhibition:* The preparation of cells, antibiotic, and 96-well plate were prepared as in (*2*). When the cefotaxime was added, clavulanic acid potassium salt (Sigma) was also added to achieve final concentrations ranging from 0-5 μg/mL.
8. *Varying initial cell density*: Culture was prepared as in (1) except culture dilution ranged from 100 to 100,000fold. Cultures were exposed to 0 or 100 μg/mL cefotaxime and time courses were measured by plate reader and the final cell density after 30 hours was recorded.
9. *Plate reader data analysis:* MATLAB (Version 9.2.0.556344, R2017a) was used to plot and characterize the time courses obtained from the plate reader (i.e. recovery time, growth rate, and change in GFP over time).
10. *Modeling:* The interaction between a β-lactam and a bacterial population expressing a β-lactamase (Bla) can be simplified to the interactions between four main components: population density, antibiotic concentration, nutrient level, and Bla concentration. To model bacteria that constitutively produce Bla and lyse due to antibiotics degrading the cell wall, we modified Tanouchi et al.’s ordinary-differential-equation model^56^ for the nondimensional dynamics of bacterial density (n), extracellular Bla concentration (B), nutrient level (S), and β-lactam concentration (A).

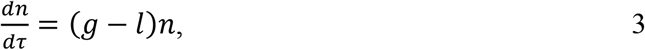

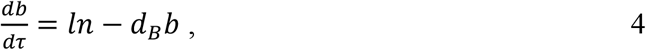

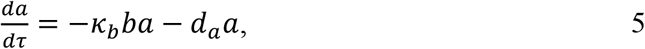

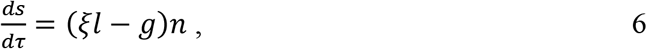

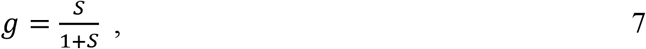

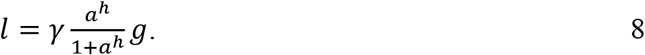 Initial conditions of *n*(0) = 0.4, *b*(0) = 0, *a*(0) = 0:300, and *s*(0) = 4 were used for all the simulations. We assume that the growth rate of cells (*g*) is limited by *s*, following the Monod kinetics. We assume the lysis rate (*l*) can reach a maximum of *γ*, depends on *a* following the Hill kinetics, and g. *d_B_* and *d_A_* are the natural decay rates of extracellular Bla and antibiotic, respectively. *κ_b_* is the rate at which Bla degrades the antibiotic. *ξ* is a weighting factor for how much nutrients is recycled upon cell lysis, and h is the Hill coefficient. See supplemental text for parameter values and full model development.

## Author contributions

H.R.M. conceived the research, designed and performed both modeling and experimental analyses, interpreted the results, and wrote the manuscript. VA assisted in model development, simulations, data interpretation, and manuscript revision. A.J. Lopatkin and A.J. Lee assisted in data interpretation and manuscript revisions. D.J.A. contributed isolates and medical perspective on how the work would clinically translate. G.B. assisted in model development and manuscript revisions. L.Y. conceived the research, assisted in research design, data interpretation, and manuscript writing. The authors declare no conflict of interest.

## Acknowledgments

Research in our lab related to the topic of this article is in part funded by the US National Institutes of Health, National Science Foundation and Army Research Office and by the David and Lucile Packard Foundation. H.R.M. acknowledges the National Science Foundation Graduate Research Fellowship Program.

## References

1 Shade A. et al. Fundamentals of microbial community resistance and resilience. (2012).

2 Nimmo, D., Mac Nally, R., Cunningham, S., Haslem, A. & Bennett, A. Vive la résistance: reviving resistance for 21st century conservation. Trends in ecology & evolution 30, 516–523 (2015).

3 Ghedini, G., Russell, B. D. & Connell, S. D. Trophic compensation reinforces resistance: herbivory absorbs the increasing effects of multiple disturbances. Ecology Letters 18, 182–187 (2015).

4 DeRose R. J. & Long J. N. Resistance and resilience: A conceptual framework for silviculture. Forest Science 60, 1205–1212 (2014).

5 Halpern C. B. Early successional pathways and the resistance and resilience of forest communities. Ecology 69, 1703–1715 (1988).

6 Connell S. D. & Ghedini G. Resisting regime-shifts: the stabilising effect of compensatory processes. Trends in ecology & evolution 30, 513–515 (2015).

7 Funk, J. L., Cleland, E. E., Suding, K. N. & Zavaleta, E. S. Restoration through reassembly: plant traits and invasion resistance. Trends in ecology & evolution 23, 695–703 (2008).

8 Holling C. S. Resilience and stability of ecological systems. Annual review of ecology and systematics 4, 1–23 (1973).

9 Todman L. et al. Defining and quantifying the resilience of responses to disturbance: a conceptual and modelling approach from soil science. Scientific reports 6 (2016).

10 Allison S. D. & Martiny J. B. Resistance, resilience, and redundancy in microbial communities. Proceedings of the National Academy of Sciences 105, 11512–11519 (2008).

11 Hodgson, D., McDonald, J. L. & Hosken, D. J. What do you mean, ‘resilient’? Trends in ecology & evolution 30, 503–506 (2015).

12 Grimm V. & Wissel C. Babel, or the ecological stability discussions: an inventory and analysis of terminology and a guide for avoiding confusion. Oecologia 109, 323–334 (1997).

13 Reller, L. B., Weinstein, M., Jorgensen, J. H. & Ferraro, M. J. Antimicrobial Susceptibility Testing: A Review of General Principles and Contemporary Practices. Clinical Infectious Diseases 49, 1749–1755, doi:10.1086/647952 (2009).

14 Pai, A., Tanouchi, Y. & You, L. Optimality and robustness in quorum sensing (QS)-mediated regulation of a costly public good enzyme. Proceedings of the National Academy of Sciences 109, 19810–19815 (2012).

15 Balaban, N. Q., Merrin, J., Chait, R., Kowalik, L. & Leibler, S. Bacterial persistence as a phenotypic switch. Science 305, 1622–1625 (2004).

16 Jorgensen J. H. & Turnidge J. D. in Manual of Clinical Microbiology, Eleventh Edition 1253–1273 (American Society of Microbiology, 2015).

17 CLSI. in Twenty-Second informational supplement NCCLS document M100-S22. National Committee for Clinical Laboratory Standards Vol. 32 1–188 (Clinical and Laboratory Standards Institute, Wayne, PA, 2012).

18 Dobrindt, U., Hochhut, B., Hentschel, U. & Hacker, J. Genomic islands in pathogenic and environmental microorganisms. Nature Reviews Microbiology 2, 414–424 (2004).

19 Neu H. C. The crisis in antibiotic resistance. Science 257, 1064–1073 (1992).

20 Alanis A. J. Resistance to antibiotics: are we in the post-antibiotic era? Archives of medical research 36, 697–705 (2005).

21 Levin B. R. & Rozen D. E. Non-inherited antibiotic resistance. Nature Reviews Microbiology 4, 556–562 (2006).

22 Levy S. B. & Marshall B. Antibacterial resistance worldwide: causes, challenges and responses. Nature medicine 10, S122–S129 (2004).

23 Livermore D. & Hawkey P. CTX-M: changing the face of ESBLs in the UK. Journal of Antimicrobial Chemotherapy 56, 451–454 (2005).

24 Rupp M. E. & Fey P. D. Extended spectrum β-lactamase (ESBL)-producing Enterobacteriaceae. Drugs 63, 353–365 (2003).

25 Ramphal R. & Ambrose P. G. Extended-spectrum β-lactamases and clinical outcomes: current data. Clinical infectious diseases 42, S164–S172 (2006).

26 Centers for Disease Control and Prevention. (ed U.S. Department of Health and Human Services) (2013).

27 Page M. G. in Antibiotic Discovery and Development 79–117 (Springer, 2012).

28 Thaden, J. T., Fowler, V. G., Sexton, D. J. & Anderson, D. J. Increasing incidence of extended-spectrum β-lactamase-producing Escherichia coli in community hospitals throughout the southeastern United States. infection control & hospital epidemiology 37, 49–54 (2016).

29 Coque, T., Baquero, F. & Canton, R. Increasing prevalence of ESBL-producing Enterobacteriaceae in Europe. Euro surveillance: bulletin Europeen sur les maladies transmissibles=European communicable disease bulletin 13, 5437–5453 (2008).

30 Villegas, M., Kattan, J., Quinteros, M. & Casellas, J. Prevalence of extended-spectrum β-lactamases in South America. Clinical Microbiology and Infection 14, 154–158 (2008).

31 Bush K. Beta-lactamase inhibitors from laboratory to clinic. Clinical microbiology reviews 1, 109–123 (1988).

32 Livermore D. M. beta-Lactamases in laboratory and clinical resistance. Clinical Microbiology Reviews 8, 557–584 (1995).

33 Lee A. J. et al. Robust, linear correlations between growth rates and β-lactam–mediated lysis rates. Proceedings of the National Academy of Sciences, 201719504 (2018).

34 Zhang, X. Y., Trame, M., Lesko, L. & Schmidt, S. Sobol sensitivity analysis: a tool to guide the development and evaluation of systems pharmacology models. CPT: Pharmacometrics & Systems Pharmacology 4, 69–79 (2015).

35 Hunter, P. A., Coleman, K., Fisher, J. & Taylor, D. In vitro synergistic properties of clavulanic acid, with ampicillin, amoxycillin and ticarcillin. Journal of Antimicrobial Chemotherapy 6, 455–470 (1980).

36 Bush K. & Jacoby G. A. Updated functional classification of β-lactamases. Antimicrobial agents and chemotherapy 54, 969–976 (2010).

37 Ball P. The clinical development and launch of amoxicillin/clavulanate for the treatment of a range of community-acquired infections. International journal of antimicrobial agents 30, 113–117 (2007).

38 Weber, D. J., Tolkoff-Rubin, N. E. & Rubin, R. H. Amoxicillin and Potassium Clavulanate: An Antibiotic Combination Mechanism of Action, Pharmacokinetics, Antimicrobial Spectrum, Clinical Efficacy and Adverse Effects. Pharmacotherapy: The Journal of Human Pharmacology and Drug Therapy 4, 122–133 (1984).

39 Lebre, P. H., De Maayer, P. & Cowan, D. A. Xerotolerant bacteria: surviving through a dry spell. Nature Reviews Microbiology (2017).

40 Colinet, H., Lee, S. F. & Hoffmann, A. Temporal expression of heat shock genes during cold stress and recovery from chill coma in adult Drosophila melanogaster. The FEBS journal 277, 174–185 (2010).

41 Lindquist S. & Craig E. The heat-shock proteins. Annual review of genetics 22, 631–677 (1988).

42 Veraart A. J. et al. Recovery rates reflect distance to a tipping point in a living system. Nature 481, 357 (2012).

43 Faassen E. J. et al. Hysteresis in an experimental phytoplankton population. Oikos 124, 1617–1623 (2015).

44 Lozupone, C. A., Stombaugh, J. I., Gordon, J. I., Jansson, J. K. & Knight, R. Diversity, stability and resilience of the human gut microbiota. Nature 489, 220–230 (2012).

45 Lewis K. Persister cells. Annual review of microbiology 64, 357–372 (2010).

46 Spratt B. G. Resistance to antibiotics mediated by target alterations. Science-AAAS-Weekly Paper Edition-including Guide to Scientific Information 264, 388–396 (1994).

47 Bugg T. D. et al. Molecular basis for vancomycin resistance in Enterococcus faecium BM4147: biosynthesis of a depsipeptide peptidoglycan precursor by vancomycin resistance proteins VanH and VanA. Biochemistry 30, 10408–10415 (1991).

48 Artemova, T., Gerardin, Y., Dudley, C., Vega, N. M. & Gore, J. Isolated cell behavior drives the evolution of antibiotic resistance. Molecular systems biology 11, 822 (2015).

49 Drawz S. M. & Bonomo R. A. Three decades of β-lactamase inhibitors. Clinical microbiology reviews 23, 160–201 (2010).

50 Watkins, R., Papp-Wallace, K. M., Drawz, S. M. & Bonomo, R. A. Novel β-lactamase inhibitors: a therapeutic hope against the scourge of multidrug resistance. Frontiers in microbiology 4, 392 (2013).

51 Letourneau, A. R., Calderwood, S. B., Hooper, D. C. & Bloom A. Combination beta-lactamase inhibitors, carbapenems, and monobactams. (2009).

52 U.S. Food and Drug Administration. (2005).

53 White A. R. et al. Augmentin (amoxicillin/clavulanate) in the treatment of community-acquired respiratory tract infection: a review of the continuing development of an innovative antimicrobial agent. Journal of Antimicrobial Chemotherapy 53, 3–20 (2004).

54 Bottenfield G. W. et al. Safety and tolerability of a new formulation (90 mg/kg/day divided every 12 h) of amoxicillin/clavulanate (Augmentin^®^) in the empiric treatment of pediatric acute otitis media caused by drug-resistant Streptococcus pneumoniae. The Pediatric infectious disease journal 17, 963–968 (1998).

55 Allen R. C. & Brown S. P. Modified antibiotic adjuvant ratios can slow and steer the evolution of resistance: co-amoxiclav as a case study. bioRxiv, 217711 (2017).

56 Tanouchi, Y., Pai, A., Buchler, N. E. & You, L. Programming stress-induced altruistic death in engineered bacteria. Molecular systems biology 8 (2012).

